# Species-aware DNA language models capture regulatory elements and their evolution

**DOI:** 10.1101/2023.01.26.525670

**Authors:** Alexander Karollus, Johannes Hingerl, Dennis Gankin, Martin Grosshauser, Kristian Klemon, Julien Gagneur

## Abstract

The rise of large-scale multi-species genome sequencing projects promises to shed new light on how genomes encode gene regulatory instructions. To this end, new algorithms are needed that can leverage conservation to capture regulatory elements while accounting for their evolution. Here we introduce species-aware DNA language models (LMs), which we trained on more than 800 species spanning over 500 million years of evolution. Investigating their ability to predict masked nucleotides from context, we show that DNA LMs distinguish transcription factor and RNA-binding protein motifs from background non-coding sequence. Owing to their flexibility, DNA LMs capture conserved regulatory elements over much further evolutionary distances than sequence alignment would allow. Remarkably, DNA LMs reconstruct motif instances bound in vivo better than unbound ones and account for the evolution of motif sequences and their positional constraints, showing that these models capture functional high-order sequence and evolutionary context. We further show that species-aware training yields improved sequence representations for endogenous and MPRA-based gene expression prediction, as well as motif discovery. Collectively, these results demonstrate that species-aware DNA language models are a powerful, flexible, and scalable tool to integrate information from large compendia of highly diverged genomes.

## Introduction

A typical eukaryotic genome contains large regions of non-coding DNA. These are not translated into proteins, but contain regulatory elements which control gene expression in response to environmental cues. Finding these regulatory elements and elucidating how their combinations and arrangements determine gene expression is a major goal of genomics research and is of great utility for synthetic biology and personalized medicine.

In the last decade, significant progress has been made towards understanding regulation of model species, particularly human and mouse, by leveraging the work of large consortia such as ENCODE^1^ or FANTOM5^2^. These groups have invested considerable resources to compile massive compendia of experiments which probe many steps of transcription control at high resolution and depth.

Nonetheless, estimates indicate that there are millions of eukaryotic species^3^. Many of these have agricultural, medical or biotechnological relevance and even those lacking direct economic importance may still hold key insights about regulatory evolution. Accordingly, it would certainly be valuable if existing approaches were extended beyond model organisms. Nevertheless, generating an ENCODE for every species is not feasible at the current level of technology. What is more in reach, however, is to sequence the genomes of all species – and this is increasingly being done^4–8^.

As the genomes of all organisms are evolutionarily related, it is possible to study regulatory elements through sequence comparison, without requiring additional experimental annotations^9^. Specifically, we expect that regulatory sequences and functional arrangements thereof are selected and thus generally more conserved than expected under neutral mutation^10^. The main method to determine whether particular nucleotides are conserved makes use of sequence alignment. Unfortunately, alignment is difficult for non-coding sequences, presumably because regulatory sequences evolve faster than coding sequences and because of a certain tolerance to the exact order, orientation, and spacing of regulatory elements in regulatory sequences^11,12^. Thus, while alignments will certainly remain valuable, their ability to integrate information across large compendia of highly diverged species is limited.

Inferring regulatory elements from genomic sequences only, without requiring further gene expression-related experimental data, i.e. labels, is reminiscent of a typical challenge in another domain, natural language processing, where vast quantities of mostly unlabelled text are available. There, masked language modeling, a type of self-supervised representation learning, presents another way to generate insights from unlabelled data^13^. To train masked language models (LMs), parts of an input sequence are hidden (masked) and the model is tasked to reconstruct them. To do so, LMs learn an internal numerical representation of words and their context, capturing the syntax and semantics of natural language. In turn, these representations can be used as features for efficiently training supervised models for many downstream tasks. This approach has alleviated scarcity of labeled data in many natural language processing predictive tasks such as translation or question answering.

In genomics, previous work has adopted masked language modeling as a method to build sequence representations for DNA^14^, although initially this was focused on single species. Only recently, multi-species datasets have been used to train large genomic language models^15–17^. These models foremost serve as a basis to build predictors of molecular phenotypes. In principle, LMs should benefit from multi-species training through the ability to leverage evolution. While first results in this direction have been very encouraging^16^, a recent analysis found that a model trained only on human sequences achieves better predictions for fruit fly enhancers compared to a multispecies model that includes the fruit fly genome^17^ – an observation which is difficult to reconcile with the 700 million years of divergence between these species^18^. Performance on downstream tasks after fine tuning is an indirect measure of the features learned by multispecies pretraining since the exact features needed to succeed at the given tasks are unknown, potentially learned during fine tuning and not investigated in these works.

Due to their focus on downstream tasks, previous works considered the actual task the language model is trained to perform – reconstructing masked nucleotides – as means to an end and do not study it. The sole exception to this trend is a recent contribution^19^, which showed that poorly reconstructed nucleotides were enriched for variants with low frequency in the *Arabidopsis thaliana* population – although the strongest variants driving this signal appear to be coding rather than regulatory. Moreover, while their model was trained on eight genomes, all evaluations were done on the *Arabidopsis thaliana* genome only, which the model was also trained on. Thus, while the authors suggest that the prediction of their language model can be considered a generalized conservation score, it is unclear whether their language model leverages between-species, rather than within-species, conservation of regulatory rules, and, more importantly, if their model accounts for changes in the code between species.

In this study, we aim to address these limitations by training masked LMs on a large number of highly evolutionarily diverged eukaryotes. Compared to previous approaches, we provide species information to our models to avoid the model having to infer the species context to account for evolution of the regulatory code. We focus on non-coding regions and explicitly evaluate whether the models have learned meaningful species-specific and shared regulatory features during training across evolution and can transfer them to unseen species. Finally, we evaluate whether sequence representations provided by the models are predictive of important molecular phenotypes, such as RNA half-life or gene expression, and encode biologically meaningful motifs.

## Results

### Using language models as an alignment-free method to capture conservation and evolution of regulatory elements

Sequence alignment is a well-known and highly effective method to study the evolution of biological sequences. It can be used to detect homologies, find conserved subsequences, and pinpoint sequence motifs. We first assessed whether sequence alignments could be a viable starting point to capture conserved regulatory sequence elements in 3’ regions across large evolutionary distances. To this end, we aligned annotated 3’ untranslated regions (UTRs) and, as control, coding sequences of *Saccharomyces cerevisiae* to the genomes of a variety of fungal species (Fig. 1A). Coding sequences could be successfully aligned even between highly diverged species. In contrast, the ability to align 3’ regions almost completely disappears beyond the *Saccharomyces* genus.

**Fig. 1:**
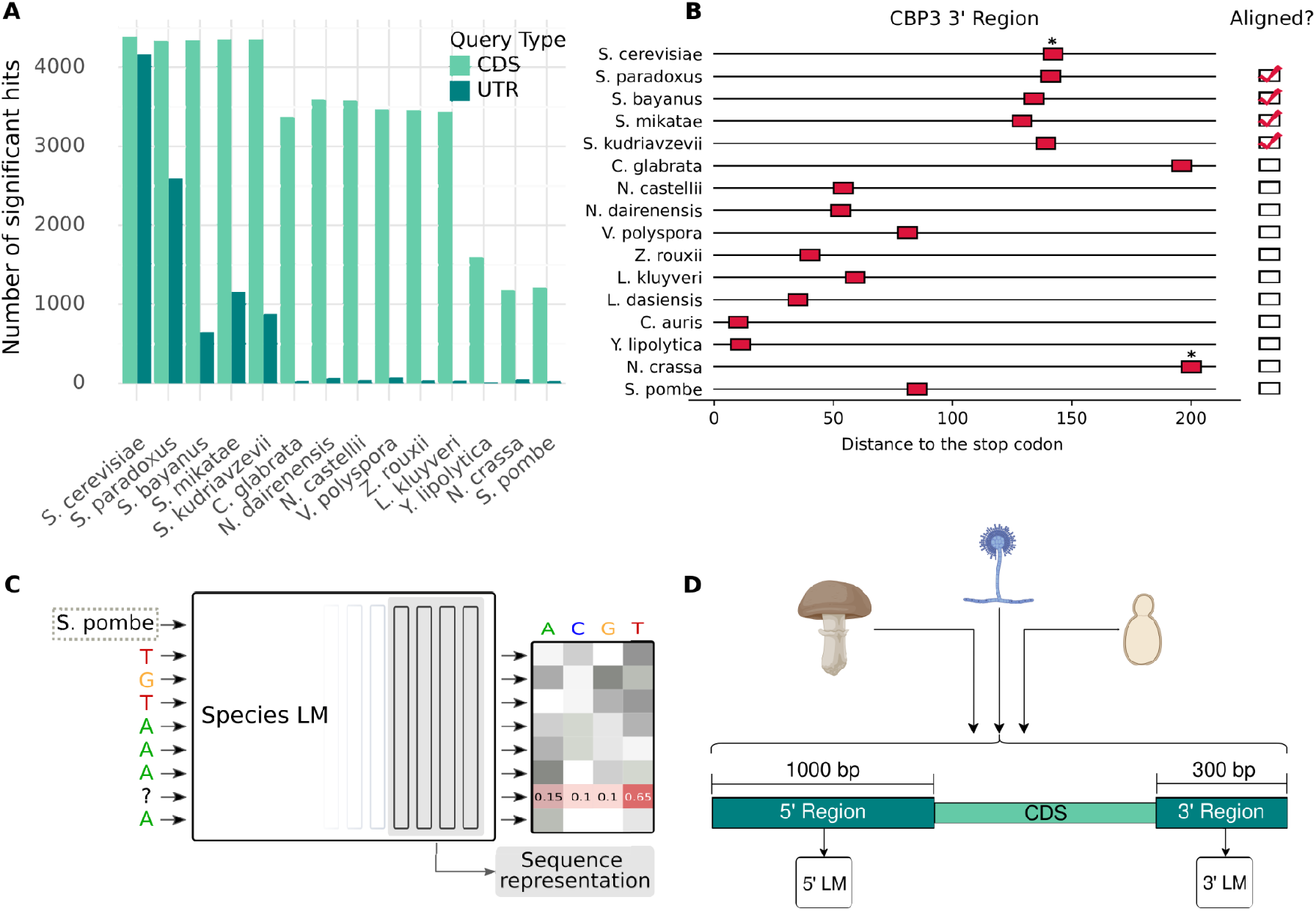
Masked language modeling can serve as an alternative for alignments, which struggle to capture conservation of regulatory elements over large evolutionary distances. **A)** BLAST hits of *S*.*cerevisiae* CDS and 3’ UTRs in other fungal species. **B)** Regions 3’ to the stop codon of CBP3 orthologues in different fungal species. Instances of Puf3 like motifs (TGTA*ATA) indicated in red, star indicates experimental evidence of Puf3 binding. It appears that Puf3 binding is conserved whereas the location of the motif is not. **C)** Masked language models are neural networks trained to reconstruct masked nucleotides from context. We illustrate this with the example of a Puf3 motif (TGTAAATA), where the second to last T has been hidden. Since this motif is highly conserved, the model may learn that, given this context, a T is most likely. For each masked nucleotide, the model returns a probability distribution over the letter A, C, G, T. Additionally, we can extract sequence representations from the model by pooling the hidden representations of the last four layers of the model. Additionally, we make the model species-aware, by providing a token denoting the species where the sequence is from originally. **D)** We train language models on hundreds of highly diverged fungi. In each genome, we locate the annotated coding sequences and we extract the non-coding sequences immediately before the start codon (5’ region) and immediately after the stop codon (3’ region). We train separate models for each region.

Nevertheless, many regulatory sequence elements of 3’ UTRs are conserved far beyond the genus boundary. The 3’ region of the cytochrome B assembly factor CBP3 illustrates this. Experiments have shown that the RNA binding protein Puf3 binds 3’ UTR of this gene in *S. cerevisiae* and the far diverged (about 500 Mya^20^) *Neurospora crassa*^21^. Moreover, we found that the Puf3 consensus binding motif can be found 3’ to the stop of the CBP3 homologue in almost all yeasts. However, we observed that the motif appears to be highly mobile, complicating alignment (Fig. 1B). This example illustrates the need and potential for alignment-free approaches to leverage conservation across large evolutionary distances.

In principle, masked language modeling should be able to leverage evolutionary information without requiring alignment because their representation of sequence is more flexible and expressive, alleviating the rigid order constraint that sequence alignments subsume. Specifically, we expect that nucleotides with regulatory function should be more conserved, within and particularly across species, and therefore easier to reconstruct when masked than remaining background non-coding sequences.

To explicitly test this, we trained masked LMs (Fig. 1C) on non-coding regions adjacent to genes extracted from a vast multispecies dataset, comprising 806 fungal species separated by more than 500 million of years of evolution. We trained distinct models for the 1,000 nucleotides 5’ of the start codon (5’ region) and the 300 nucleotides 3’ of the stop codon (3’ region) of all annotated coding sequences (Fig. 1D). The 5’ region typically contains the 5’UTR and the promoter of the gene^22^. The 3’ region typically contains the 3’ UTR^23^. Hence, we expect these regions to be enriched for non-coding sequences and to capture both transcriptional regulatory elements as well as post-transcriptional regulatory elements involved in mRNA stability and translation. Importantly, no annotation of UTR, promoters, nor transcription start site or polyA site were provided to the model. Models were trained with (species LM) and without species labels (agnostic LM). We focused on fungi, since many fungal genomes are available, they evolve quickly and their transcriptional control generally makes less use of extreme long-range interactions which are difficult to model with current approaches. We held out the entire *Saccharomyces* genus to test the generalization performance of our model in the well-studied species *Saccharomyces cerevisiae*.

### Language models reconstruct known binding motifs in an unseen species

Binding motifs, which represent the sequence specificities of DNA and RNA binding proteins (RBP), are considered to be the “atomic units of gene expression”^24^. Accordingly, to verify that LMs can capture aspects of the regulatory code in an alignment-free manner, we first needed to verify that they capture important known motifs.

To test this, we analyzed to what extent our 3’ and 5’ LMs could reconstruct nucleotides in the held-out species *S. cerevisiae*. We compared the reconstruction obtained by our LMs with a number of baselines, including approaches based on k-mer frequencies and alignment of species beyond the genus. We also computed the reconstruction achieved by aligning *S. cerevisiae* with other *Saccharomyces* species, which can be regarded as an estimate of the upper range of achievable reconstruction.

We found that all approaches – except the alignment of close species – perform similarly when we compute the reconstruction accuracy over all nucleotides (Fig. 2A and SFig. 1). This changes drastically when considering the reconstruction of known motifs. Instances matching the Puf3 consensus motif, for example, are reconstructed almost as well by the species-aware 3’ model as they are by the alignment of close species (Fig. 2A), strongly outperforming the alignment with far species and other baselines. We observe similar results for other RBP motifs, as well as the consensus motifs of a number of transcription factors (TF) in the 5’ region (SFig. 1). Remarkably, by applying Modisco clustering^25^ on all nucleotides weighted by their information content as computed by our LMs, we were able to recover some of these motifs *de novo* (Fig. 2B, SFig. 2).

**Fig. 2:**
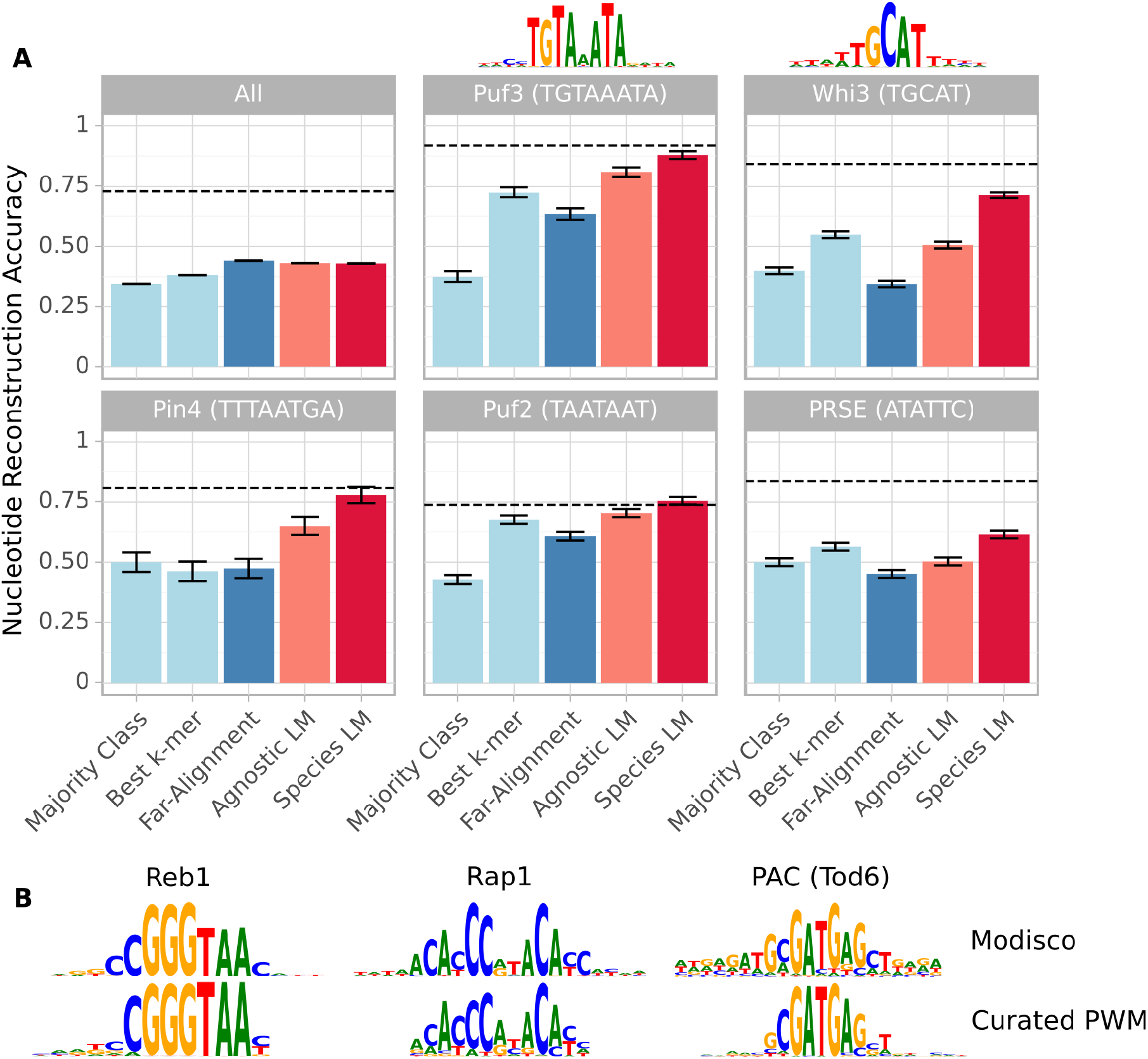
Language models reconstruct likely regulatory sequences in a held-out species and recover known binding motifs. **A)** Reconstruction accuracy for nucleotides within instances of RNA-binding protein consensus motifs and across all nucleotides in *S. cerevisiae* 3’ UTR sequences (those longer than 300 bp have been truncated). We compare the agnostic and species 3’ LM to a variety of baselines. Dashed line represents the accuracy achieved by the intra-genus alignment. For Puf3 and Whi3, Modisco clustering on the species LM reconstructions recovers the motif (depicted above the respective plots). **B)** A sample of known transcription factor motifs recovered by applying Modisco clustering to the 5’ species LM reconstructions, (manually) matched to the respective high-confidence PWM from the YeTFaSCo database.

For many motifs we tested, the species-aware language models reconstructed slightly better than their already strong agnostic counterparts. In sum, our analysis clearly demonstrates that language models trained on a large compendium of highly-diverged genomes are (1) able to learn conserved regulatory elements and (2) able to transfer this knowledge to unseen species.

### Context-sensitive reconstruction of motifs is predictive of in vivo binding

It has been shown for many genomes that only a fraction of the instances of a particular motif is functional, i.e. binds the respective protein in vivo^26–28^. This is because, among other reasons, binding also depends on the context of the motif. Therefore, we next sought to evaluate whether our models can recognize these relationships.

An important piece of context for many motifs is their position with respect to certain genomic landmarks. One example of this is that RBP motifs can only be functional if they are located in a transcribed region. Another example is that in yeast TF binding sites tend to be located relatively close to the transcription start site (TSS)^29^. As a first test of whether our LMs are capable of locating these genomic landmarks – despite their location not having been indicated during training – we computed the actual and predicted nucleotide biases as a function of the distance to the TSS (imputed using CAGE data^30^) and the distance to the end of the 3’ UTR^23^. In both cases, we observed that the LMs track local changes in nucleotide biases (Fig. 3A, SFig. 3).

**Fig. 3:**
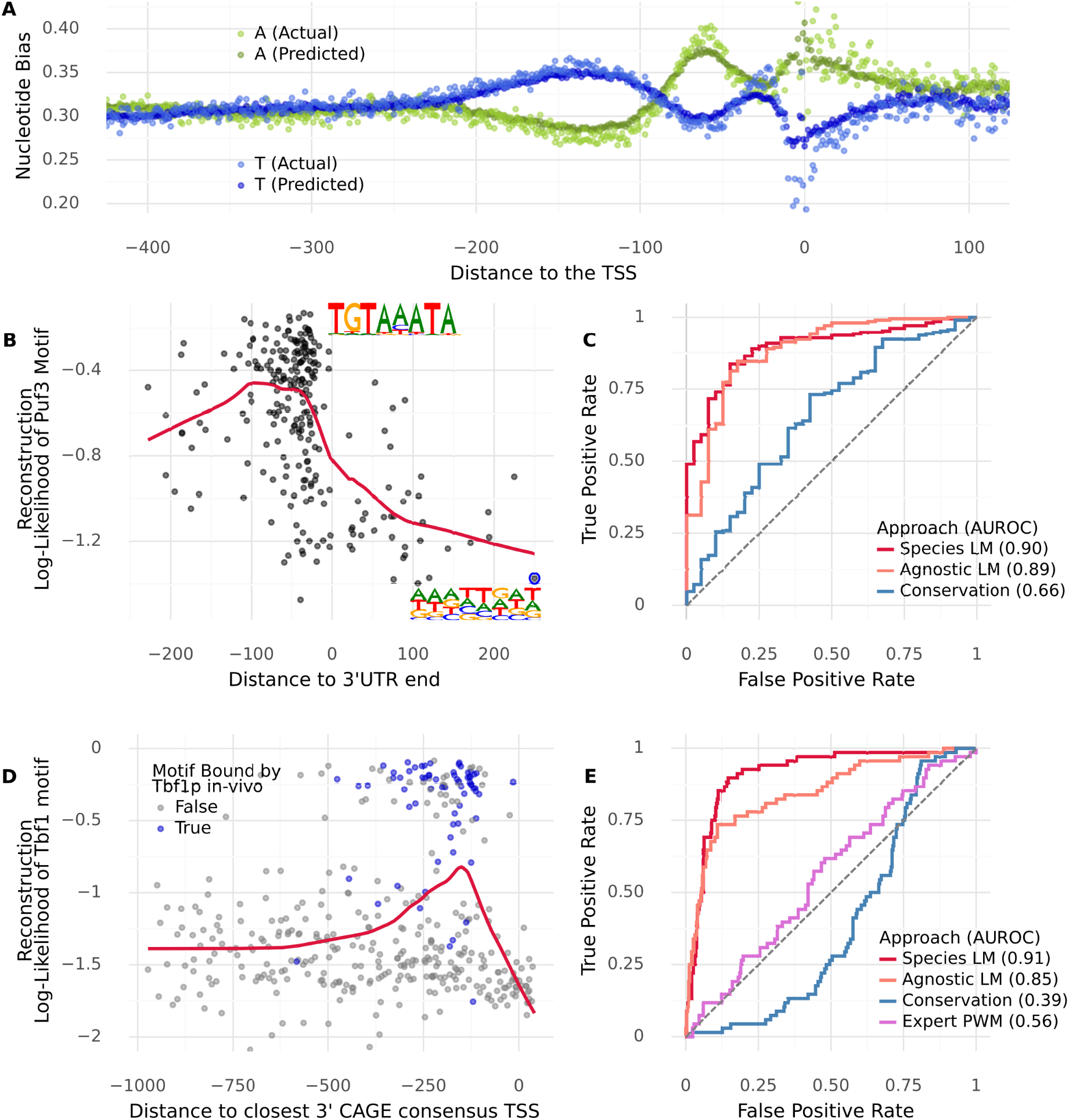
Reconstruction of motifs depends on the context and predicts whether a motif instance will be bound in vivo. **A)** Actual and predicted (by the 5’ species LM) nucleotide biases as function of the distance to the TSS (imputed using CAGE data). The model keeps track of local variations in nucleotide biases. **B)** Reconstruction fidelity (log-likelihood of the observed nucleotides according to the 3’ Species LM) of instances of the Puf3 motif (TGTAAATA), as a function of the distance to the end of the annotated 3’ UTR. The predictions of the model for masked nucleotides are indicated for two instances (blue points). Reconstruction fidelity is notably degraded beyond the 3’ UTR end. **C)** ROC curve evaluating to what extent the reconstruction fidelity of our 3’ LMs, as well as the phastCons conservation score, can serve as a predictor of whether a Puf3 motif instance is within or beyond the 3’ UTR boundary. The LMs greatly outperform the conservation score. **D)** Reconstruction fidelity (log-likelihood of the observed nucleotides according to the 5’ Species LM) of instances of the Tbf1 consensus motif (ARCCCTA), as a function of the distance to the closest 3’ TSS (imputed using CAGE data). Blue indicates that the motif instance was bound in vivo according to Chip-exo data. Motif instances that are around −100 to −250 nt to the TSS are better reconstructed than those further away or in the 5’ UTR. **E)** ROC curve evaluating to what extent the reconstruction fidelity of our 5’ LMs, as well as the phastCons conservation score and an expert-curated PWM, can serve as a predictor of whether a Tbf1 motif instance is bound in vivo. The LMs again greatly outperform the alternative methods.

To explicitly test whether the 3’ LM locates and accounts for genomic landmarks when reconstructing motifs, we compared the reconstruction of instances of the Puf3 consensus motif located within annotated 3’ UTR regions to those which are located beyond the 3’ UTR yet still within 300 bp of the stop codon. We find that motif instances within the annotated 3’ UTR are reconstructed significantly better (Fig. 3B-C), consistent with the function of RBPs. In contrast, the phastCons^31^ score, an alignment-based measure of conservation, appeared to be a poor predictor of whether a Puf3 site is within a 3’ UTR or not. We repeated this analysis for other 3’ UTR motifs, finding similar results (SFig. 4A,B). For a TF, we expected this relationship to be reversed, and indeed the E-box motif was reconstructed better when found outside of 3’ UTR regions (SFig. 4C).

**Fig. 4:**
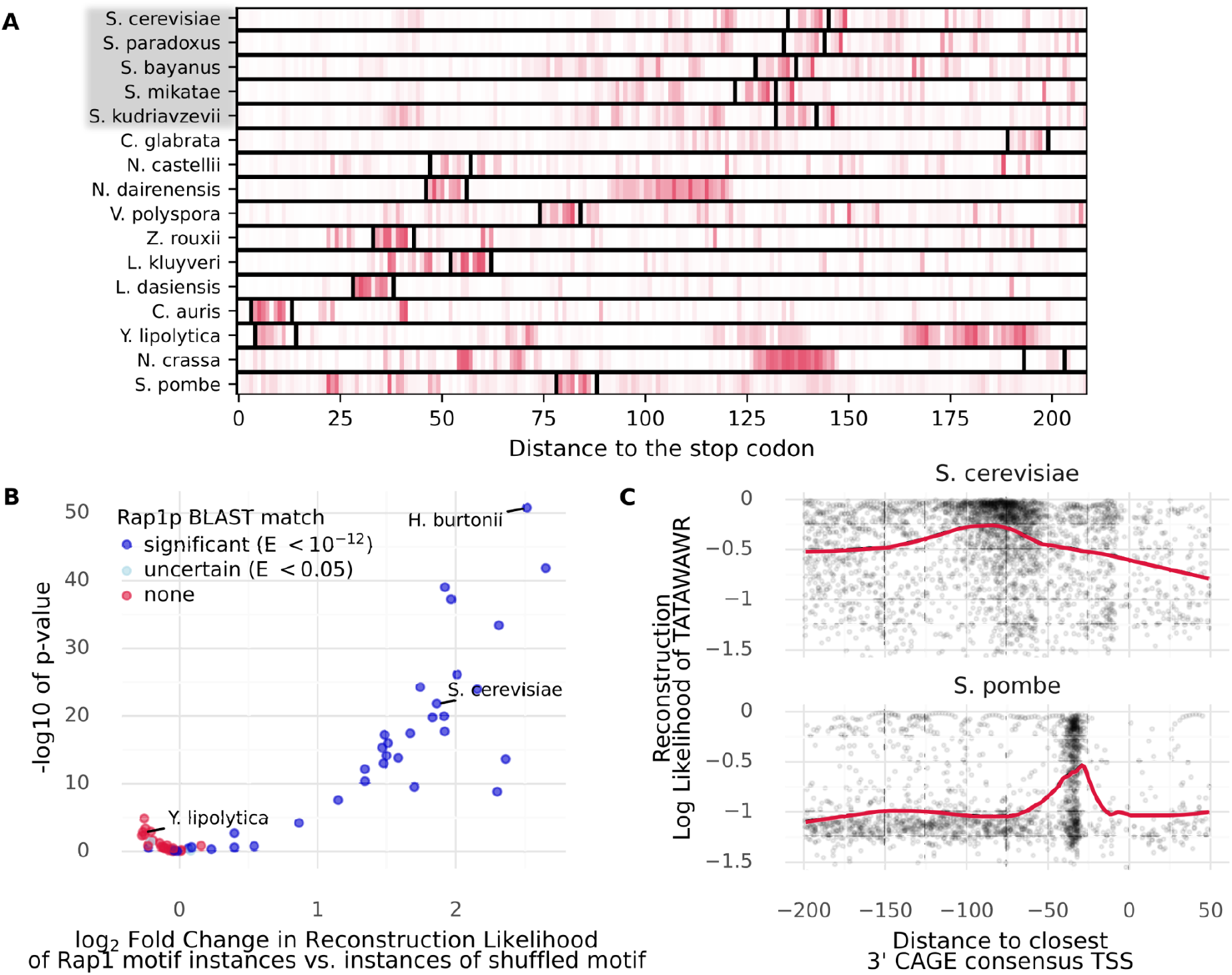
LMs can trace the movement and disappearance of motifs across species and they account for the evolution of the transcription initiation mechanism. **A)** We applied the 3’ species LM to the 3’ region of CBP3 homologues in a number of fungal species (compare Fig. 1A). Darker color indicates that the model assigns a higher probability to the correct nucleotide at that position. In most species, the Puf3 motif instance (delineated with black bars) is notably reconstructed better than the background. Species with gray background were held out during LM training. **B)** We computed the reconstruction fidelity (log-likelihood) achieved by the species 5’ LM for the *S. cerevisiae* consensus Rap1 motif (CAYCCRTACAY) instances and for instances matching shuffled versions of this motif in 60 fungal species. The difference in reconstruction between the true and shuffled motif instances, expressed as log_2_ fold change, is plotted against the -log_10_ p-value of this difference, computed using a Mann-Whitney-U test. We observe that in species that have no BLAST match to *S. cerevisiae* Rap1p, the reconstruction fidelity of the *S. cerevisiae* Rap1 motif is generally not much better than that of shuffled versions thereof, indicating that the model correctly accounts for species-context when reconstructing motifs. **C)** Reconstruction fidelity (log-likelihood of the observed nucleotides according to the 5’ Species LM) of instances of the TATA-box (TATAWAWR), as a function of the distance to the closest 3’ TSS (imputed using CAGE data). Positive values indicate that the TATA-box instance is located in the 5’UTR. We observe that in *S. pombe*, the TATA-box is best reconstructed when located ca. −30 bp to the TSS. In *S. cerevisiae*, which uses a scanning mechanism to initiate transcription and therefore allows more flexible positioning of the TATA, the model reconstructs TATA well overall, but somewhat better when located 50 to 120 bp 5’ from the TSS.

Having shown that the models did not simply learn a mere lexicon of over-represented motifs, but instead seemed to account for the context in which motif instances occur, we next asked whether the reconstruction fidelity could predict whether a motif is bound in vivo. To this end, we compared the reconstruction of Puf3 motif instances located 3’ of genes that have been experimentally verified to bind Puf3p^32^ to those without verified binding. Strikingly, we found that the reconstruction fidelity serves as a predictor of whether a gene containing a Puf3 consensus motif is bound by Puf3p – despite our models never having been exposed to binding data (SFig. 5).

We repeated a similar analysis for Tbf1 (Fig. 3D) and a variety of other transcription factors (SFig. 6). Specifically, we evaluated the reconstruction of the consensus binding motifs of these TFs as a function of their distance to the closest upstream TSS. We observed that reconstruction improved when the motifs were located at a biologically plausible distance^29^ from the TSS. Moreover, the reconstruction fidelity is highly predictive of in vivo binding to motif instances as measured by Chip-Exo^33^, outperforming the phastCons score and, in some cases, expert-curated PWMs constructed using binding data (Fig. 3E, SFig. 6)^34^.

Distinct motifs have been established to exhibit associations with specific groups of co-regulated genes. An example in *S. cerevisiae* is the Rap1 motif, which is found primarily in the promoters of ribosomal protein genes^35,36^. Accordingly, we find that instances of the Rap1 consensus motifs tend to be better reconstructed if they are found within 1kb 5’ of a ribosomal protein gene (SFig. 7A). In other words, the reconstruction of the Rap1 motif serves as an indirect predictor of whether a gene belongs to the ribosomal protein module. We performed a similar analysis for the RRPE motif, which is primarily found near genes involved in ribosome biogenesis, and obtained similar results (SFig. 7B).

Overall, this analysis demonstrates that LMs do not just learn a lexicon of conserved motifs, but additionally pick up on correlations between the motifs and their context which are predictive of whether motif instances are bound in vivo. Notably, this is achieved purely from genomic sequences without requiring any additional experimental data during training. Finally, the ability to outperform the phastCons conservation score shows the advantages of an alignment-free approach.

### Language Models account for changes in the regulatory code between species

Our previous analyses have focused on the held-out species *S. cerevisiae*. However, one of the main use cases we envision for genomic LMs is to serve as a method to explore understudied species. We thus analyzed how the LMs perform when evaluated across fungal species.

In Figure 1B, we showed that the Puf3 motif, but not its location, is conserved in the 3’ regions of CBP3 homologues. We applied our 3’ model to these sequences and found that, in most species, the Puf3 motif tends to be well reconstructed compared to the background (Fig. 4A). We next applied Modisco clustering to the predictions of the model on the 3’ regions of all CBP3 homologues in our dataset. We recovered the Puf3 motif, as well as two versions of the Puf4 motif (SFig. 8A)^37^. This indicates that our method allows motif discovery not just across genes in one organism, but also for individual genes across organisms.

Over evolutionary timescales, certain motifs may either change drastically or fully disappear, particularly if the binding protein evolves or is lost. As an example, we considered Rap1p, a conserved protein known to control telomere length^38^. However, as noted previously, Rap1p additionally acts as a regulator of ribosomal protein expression in certain parts of the yeast lineage, a change that is associated with the acquisition of a transactivation domain by Rap1p^36^. To determine whether our models can reflect such changes, we analyzed the reconstruction of the *S. cerevisiae* Rap1 motif in 60 fungal species. Specifically, we computed for each species the difference in reconstruction of instances of the consensus motif and instances that correspond to shuffled versions thereof. This procedure controls for GC content and general differences in reconstruction fidelity between species. We found that in species close to *S. cerevisiae*, the motif instances are reconstructed significantly better than the shuffled instances (Fig. 4B). By contrast, in species where we cannot find a significant BLAST match to *S. cerevisiae* Rap1p we observed no such enrichment. An example of this is *Y. lipolytica*, a species known to not have a Rap1p homologue^39^. We performed a similar analysis for two other motifs, finding consistent results (SFig. 8B-C).

In addition to motif evolution, the proper context of motifs can change as well. A famous example is the positional constraint of the TATA box. In most eukaryotes including the *Schizosaccharomyces* yeasts, the TATA box is preferentially located about 30 bp 5’ from the TSS^22^. Budding yeasts, however, use a scanning mechanism to initiate transcription and therefore the position of the TATA box in these species is more flexible, but usually located between 50 and 120 bp 5’ from the TSS^22^. To verify whether our model correctly recapitulates these constraints, we located all instances of TATAWAWR in the respective species. We then assessed how well the model reconstructs these nucleotides as a function of the distance to the closest upstream TSS^30^. We found that in *S. pombe*, reconstruction fidelity of the TATA box notably peaks around 30 bp 5’ to the TSS, whereas instances located further or beyond the TSS are generally reconstructed poorly (Fig. 4C). In *S. cerevisiae*, on the other hand, TATA boxes were generally well reconstructed (likely also reflecting the AT-bias of the budding yeasts), but we observed a peak in the region 120 to 50 bp 5’ of the TSS. Thus, the model applies specific constraints when reconstructing motifs in a way that reflects the evolution of the initiation code.

In conclusion, we find considerable evidence that LMs can account for changes in the regulatory code and learn meaningful motif-context relationships in a species-specific manner.

### Species-aware language model representations encode biologically meaningful features and are directly predictive of many molecular phenotypes

While our previous analyses demonstrated that the reconstructions of any LM are already very informative and could potentially be used to explore regulation in understudied species, we can also extract the learned sequence representations and leverage them to predict gene-expression related traits in a supervised fashion. We note that this makes most sense for tasks that are data constrained, such as gene-level measures of expression or half-life where there can only be as many data points as there are unique genes/transcripts. Accordingly, to test the predictive power of the representations themselves, we selected several gene-level omics assays, including RNA half-life measurements in *S. cerevisiae* and *S. pombe*, RNA-seq based gene expression in *S. cerevisiae* and microarray measures of condition-specific gene expression for a number of yeast species^41–45^. We additionally included three reporter assays testing 3’ sequences^40,46^ and promoter sequences in isolation^47^.

We then used our masked language models to generate sequence representations for the different sequences assayed in the respective experiments (Methods). We trained linear models in cross-validation using LM representations as input for different tasks. We used linear models specifically to ensure that the predictive power derives mostly from the LM and not from a fine-tuning procedure or a heavily engineered nonlinear fitting with many tunable hyperparameters. As a baseline for what can be achieved using “naive” sequence representations, we also trained linear models on k-mer counts. Furthermore, if available, we compared against state of the art.

We found that LM sequence representations outperform simple sequence representations based on k-mer counts across all tasks, a finding consistent with previous work^16^. Remarkably, the species LM performed best on all tasks (Fig. 5A, STable 1). On a task where most of the signal comes from the coding sequence, it performed on par (better, but not significantly) with expert hand-crafted features (Cheng et al.^41^) to predict mRNA half-life. Furthermore, simple ridge regressions trained on species-aware representations significantly outperformed hyperparameter-optimized deep neural networks (Zrimec et al.^44^) on gene expression prediction from non-coding sequence. This clearly shows that species LMs learn sequence representations rich in information without requiring labeled data.

**Fig. 5:**
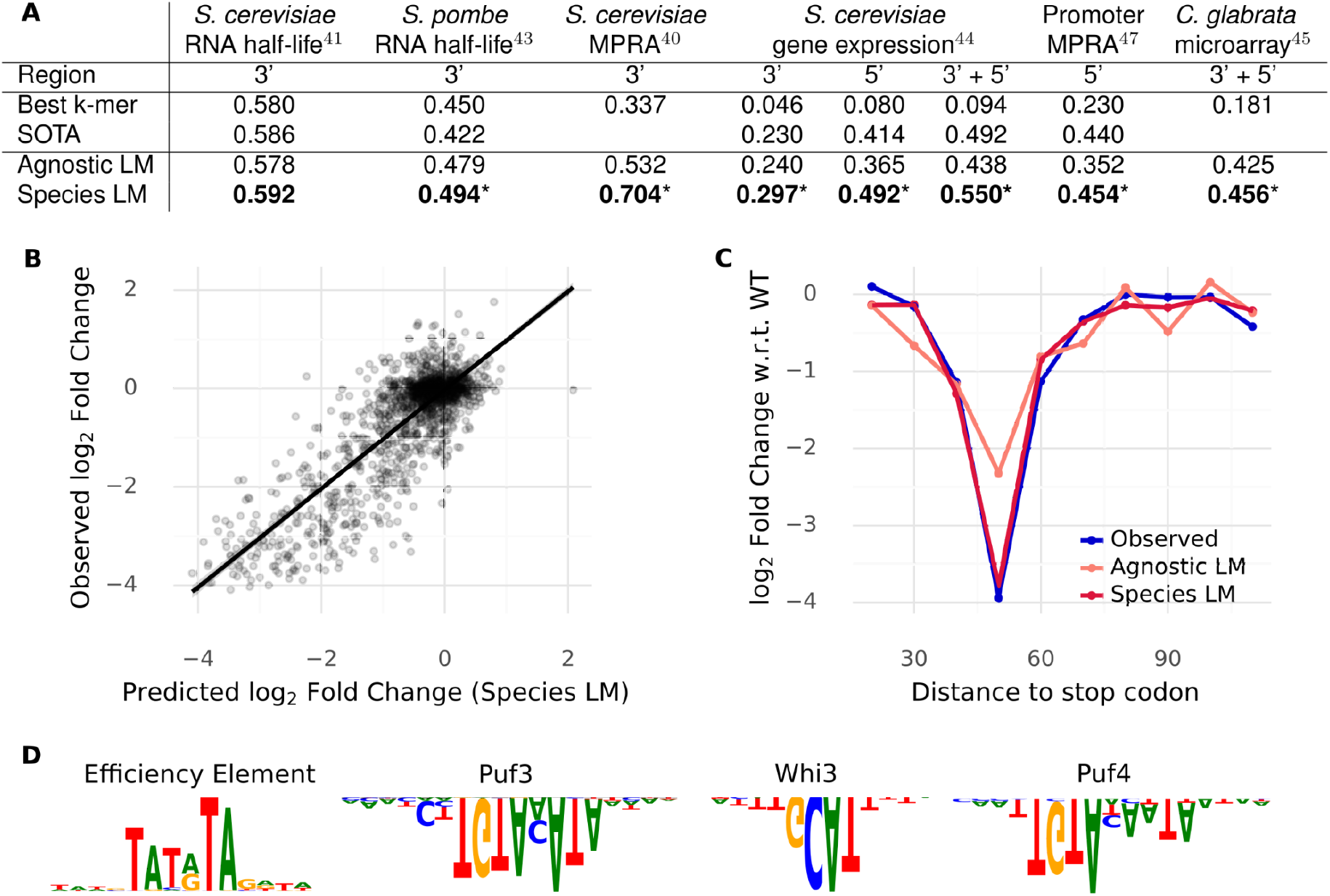
Sequence representations of the species LM outperform other methods on a variety of downstream tasks. **A)** Performance (R^2^) of linear models trained on embeddings from language models compared to state-of-the-art models and k-mer count regressions, where the best k from {3, 4, 5} is shown. Star indicates that the Species LM significantly (*P* < 0.05) outperforms the second best. **B)** Effect of mutation of 3’ sequences on expression. Observed log_2_ fold changes, as measured in Shalem et al.^40^ are well predicted by the species LM representation. **C)** Observed and predicted effect of mutation on expression as a function of distance to the stop codon for the YDR131C 3’ sequence. **D)** Motifs recovered through in-silico mutagenesis followed by Modisco clustering on our linear model for the *S. cerevisiae* half-life task. We recover (2 of 4) motifs found by Cheng et al.^41^: the Puf3 motif and the Whi3 motif. Additionally, we find two motifs not found by this previous analysis, the Puf4 motif and the efficiency element, both of which have known effects on RNA stability.

One of the datasets we considered, the Shalem et al. 3’ MPRA^40^, used a tiled mutagenesis design to measure the effect of individual subsequences of native 3’ sequences on expression. Here, the species LM performs extremely well, outperforming the agnostic model by 20 percentage points of explained variation (Fig. 5B,C). The results on this controlled experiment demonstrate that supervised models trained on species-LM sequence representations can capture causal determinants of expression found in 3’ UTR sequences.

While LM representations prove advantageous on species included in the dataset (*S. pombe*), we emphasize that they generalize to unseen species (*S. cerevisiae*). To ensure that LM performance is independent of dataset composition, and to ascertain that training on more species is beneficial, we repeated these analyses using different models pre-trained on different sets of species. We found that for all tasks, training on more species in a species-aware fashion led to better predictions (STable 2).

However, in genomics, models are often used not primarily for prediction but rather as a method to discover motifs or mechanisms in large datasets. To verify that supervised learning models trained on LM sequence representations are suitable for motif discovery, we applied a standard model interpretation workflow (in-silico mutagenesis, followed by Modisco clustering, Methods) to the *S. cerevisiae* mRNA half-life data. In this way, we recovered two out of the four motifs originally found by Cheng et al.^41^: the Puf3 and the Whi3 motif – both of which our model correctly predicts as being destabilizing (Fig. 5D). We additionally found one more destabilizing element, namely the Puf4 motif^48^ and one stabilizing element, the efficiency motif, which was shown to have large positive effects on expression in the Shalem mutagenesis reporter assay^40^ – where we also recovered this motif using the same technique. In sum, applying standard interpretation techniques with minimal manual tuning to our embedding-based approach yielded biologically meaningful sequence features.

## Discussion

In this study, we trained language models on the genomes of hundreds of fungal species, spanning more than 500 million years of evolution. We specifically directed our attention to non-coding regions, examining the models’ ability to acquire meaningful species-specific and shared regulatory attributes when trained on the genomes of many species. To our knowledge, we are the first to show that LMs are able to transfer these attributes to unseen species.

Through analysis of the masked nucleotide reconstructions provided by the models, we have demonstrated that they not only can preferentially reconstruct motifs, but they do so in a way that is sensitive to context. As a result, the reconstruction fidelity can serve as a predictor of whether a particular instance of a consensus motif will be bound in vivo. This suggests that these models could be used to discover candidate high-affinity regulatory elements in species where no binding data is available.

We have further illustrated that the models better reconstruct RBP binding sites if they are located within annotated 3’ UTRs and exhibit improved reconstruction of TATA-box instances if they are placed at a distance to the TSS that is appropriate for the given species. This is remarkable, as we indicated neither the TSS site nor the polyadenylation site during training. This suggests that the models can infer the location of these genomic landmarks. Consequently, LMs may prove useful for accurate genome annotation of understudied species.

Altogether, these analyses further indicate that the reconstruction of masked nucleotides, rather than just a means to an end, can be very informative by itself. We have focused on verifying that the models capture known features of the regulatory code, but similar techniques could potentially reveal novel associations between regulatory elements and their context.

Additionally, we have shown that providing species information to DNA LMs greatly improves their internal numerical representations of regulatory sequences. Strikingly, these representations, when used as input for simple linear models, achieve state-of-the-art predictive accuracy on a wide variety of tasks such as prediction of RNA abundance, condition-specific RNA expression, or mRNA half-life. We note that this approach requires neither retraining the whole model nor engineering a complex nonlinear downstream predictor. Despite its simplicity, we show that this procedure nevertheless recovers biologically meaningful motifs. Thus, LMs do not only learn predictive sequence representations but can also serve as a tool for biological discovery. As species awareness provides significant improvements at practically no cost (one additional token), we think that integrating a species representation will be useful for almost all DNA LMs.

This notwithstanding, the language modeling approach does have two important drawbacks. Firstly, it is computationally costly, particularly for longer sequence contexts. We initially experimented with state space models^49,50^, which scaled better to longer sequences. However, we ultimately adopted the standard DNABERT transformer architecture for simplicity, as our main goal was to explore the suitability of language models for multi-species modeling rather than a comprehensive evaluation of model architectures. Scaling language models to mammals with long-range regulatory interactions while maintaining single nucleotide resolution will require architectural improvements^51^.

While we showed that LMs generalize across highly diverged species, our analysis focused on a single kingdom. It therefore necessarily falls short of demonstrating that training on the entire eukaryotic tree of life – from protists to blue whales – could further improve language models. It is conceivable that at a certain point regulatory mechanisms diverge so fundamentally that proper generalization is no longer possible. Additionally, massive differences in genome size may require careful dataset curation. On the other hand, we did not evaluate whether LMs would benefit from large collections of very similar species, such as the recently released 233 primate genomes^6^. Lastly, we also must note that most existing genome collections do not represent a random sample of the true phylogeny but tend to oversample species with medical or biotechnological relevance while undersampling non-western species and those which are hard to isolate^52^. Systematically studying how dataset composition influences what the model learns and how well it performs in different target species is an important avenue for future work.

## Materials and Methods

### Genome data

We obtained 1,500 fungal genomes, comprising 806 different species, from the Ensembl fungi 53 database^53^. For each annotated protein-coding gene in each genome, we extracted 300 base pairs 3’ to the stop codon of genes and 1000 bases 5’ to the start codon. While the actual transcribed 3’ untranslated regions (3’ UTR) vary in length, we expect that in most cases 300 bp will be sufficient to include the entire 3’ UTR^23^. Equally, in most species 1kb should be sufficient to cover the 5’ UTR and promoter – and scaling beyond this length becomes computationally infeasible for the modeling approach taken here. Overall our train set included in the order of 13 million sequences, meaning that the 3’ models are trained on 3.9 billion nucleotides, whereas the 5’ models were trained on 13 billion nucleotides.

As a test set, we used the widely studied species *Saccharomyces cerevisiae*. To prevent data leakage from closely related species, the train set excludes the entire *Saccharomyces* genus.

### Sequence alignment

Sequence alignment of annotated *Saccharomyces cerevisiae* CDS and 3’ UTR sequences was performed using discontiguous megablast, which was specifically designed for cross-species alignment, version 2.13 with default parameters^54^. Sequence alignment for *S. cerevisiae* proteins was performed using tBlastn, with default parameters.

### Masked language modeling

We performed masked language modeling on the fungi 3’ and 5’ regions. Specifically, we randomly masked nucleotides in each sequence and trained DNABERT models^14^ to reconstruct these from context, so as to minimize cross-entropy.

For DNABERT, nucleotides were tokenized into overlapping six-mers before being passed to the model. In this context, we mask spans of overlapping 6-mers so that 80% of the nucleotides selected for masking were masked, 10% percent were randomly mutated, and the remaining 10% were left unchanged^13^. The model consists of 12 transformer encoder blocks and has around 90M parameters. We employed Flash-attention^55^ as a fast exact attention implementation. Models were trained for 200,000 steps using a batch size of 2048 using the Adam optimizer^56^. The learning rate was warmed up to 4e-4 during the first 10,000 steps and then linearly decayed to 0 until training terminates. We increased the masking ratio after 100,000 steps from 15% to 20%. Note that these hyperparameters are the same as those used in the DNABERT paper.

To make the model species aware, the species label corresponding to each region was provided as an additional input token and prepended to the sequence. At the beginning of training the species token embedding was randomly initialized and learned during training. For purposes of comparison, we also trained a species-agnostic version of the model, which does not receive the species label.

As the test species was held out from the training set, the species LM could not be provided with the matching species token. To allow it to predict anyways, we provided the model with a proxy species token of a closely related species. For the 3’ species LM we used *C. glabrata*, for the 5’ species LM we used *K. africana* for all analyses. These proxy species were selected based on overall reconstruction achieved.

We trained ablations of the 3’ models where we varied the dataset composition. The substantial computational cost of training the model prevented us from exploring more 5’ models.

### Nucleotide reconstruction

We computed reconstruction predictions for each position in the 3’ and 5’ sequences of the test species *S. cerevisiae* using the respective species and agnostic LMs. For this, we masked each nucleotide individually by masking the span of six overlapping 6-mers that contains this nucleotide. We then averaged the prediction for this nucleotide over the overlapping six-mers, to obtain one probability distribution per position.

To fit the k-mer-models, we tabulated the frequencies of nucleotides conditional on the identity of the (k-1)/2 flanking nucleotides, where k is an odd number {7,…,13} across our training dataset. To reconstruct masked nucleotides, we extracted the (k-1)/2 flanking nucleotides on either side of this masked position and then calculated the probability of each nucleotide accordingly.

To reconstruct using alignment, we first downloaded the seven yeast alignment^31^ and found the aligned position in the other species for each position in *S. cerevisiae*. We then computed the frequency over the nucleotides in the aligned positions to obtain a distribution. For the far-alignment we used the species *N. castellii* and *L. kluyveri*. For the intra-genus alignment, we used the *Saccharomyces* species.

To evaluate reconstruction for a particular motif, we used regex search to find positions in the 3’ or 5’ sequences matching the motif consensus. We allowed overlapping matches. When 5’ sequences overlap (which can occur, as they are 1kb long), we do not double count matches, but instead keep only the match in the sequence where it is located closest to the upstream gene.

Many consensus motifs we used in our analyses have degenerate positions. When we computed metrics such as reconstruction accuracy or log-likelihood on these motifs, we only take into account the non-degenerate positions, as otherwise the metrics are inflated.

For all analyses where we use reconstruction fidelity to predict some kind of outcome, including whether the motif is in the 3’ UTR, whether it is bound and whether it is reconstructed better than a shuffled version, we always use the reconstruction log-likelihood as the metric. This is mainly because accuracy, while easier to interpret, is also more likely to produce ties, as it is fundamentally a binary metric.

### 3’ UTR and TSS annotations

We extracted 3’ UTR annotations for *S. cerevisiae* from Cheng et al.^41^, who derived them from Pelechano et al.^23^, and matched them to the 3’ sequences. Note that these annotations were only available for 4388 genes.

To get TSS annotations, we gathered the consensus CAGE clusters for *S. cerevisiae* in YPD from YeastTSS^30^. We matched the locations of these consensus TSS sites with the 5’ sequences. We then matched motif instances to their closest 3’ TSS (or to the closest 5’ TSS site, if there was no 3’ TSS site between the motif and the start codon of the gene).

### Binding data

To test whether the reconstruction fidelity is predictive of Puf3 binding, we collected all instances of the Puf3 consensus motifs 3’ (i.e. within 300bp 3’ of the stop codon) of *S. cerevisiae genes*. We considered as a positive set all instances of the Puf3 motif in genes with experimental evidence of Puf3p binding their 3’ UTR^32^. We resorted to this as the data does not indicate the exact binding site, just that the 3’ UTR was bound. The negative set comprised the remaining instances. We then computed the reconstruction log-likelihood for all motifs of interest. PhastCons conservation scores^31^ for *S. cerevisiae* were downloaded from UCSC and extracted for regions of interest.

For transcription factors, we matched instances of the consensus motif with Chip-exo peaks^33^. Specifically, the data indicates the center of Chip-exo peaks. We extend this by 10bp to either side and consider that an instance of the motif was bound if it overlapped the extended peak.

### Downstream tasks

To evaluate the predictiveness of LM sequence representations, we considered the following tasks. Sun et al.^42^ measured half-life for 4388 *S. cerevisiae* mRNAs using nonperturbing metabolic RNA labeling. Cheng et al.^41^ used this data to build a quantitative model to predict mRNA half-life from sequence, using handcrafted features from the coding sequence, 5’ and 3’ UTR – which to our knowledge is the state-of-the-art model of mRNA stability in yeast. Eser et al.^43^ measured mRNA half-life in *S. pombe*. In the Shalem et al.^40^ MPRA, the expression of a fixed reporter gene was measured when combined with different 3’ sequences. These include genomic 3’ sequences as well as mutated versions of these sequences where mutations were performed in such a way as to tile the sequence. Zrimec et al.^44^ aggregated over 20,000 RNA-Seq experiments and built a convolutional neural network to predict the variation in mRNA abundance between genes from sequence. To our knowledge, this represents the state of the art model for predicting endogenous gene expression in *S. cerevisiae*. Zrimec et al. also consider two additional tasks to evaluate the generalization of their model, which we adopt. Firstly, Keren et al.^47^ measured, using a fluorescence reporter assay, the expression of a fixed reporter gene when combined with different endogenous *S. cerevisiae* promoter sequences. Yamanishi et al.^46^ also used a fluorescence reporter assay to measure the impact of different *S. cerevisiae* terminator sequences. Finally, to have data from non-model species, we obtain microarray data measuring condition specific gene expression in different stages of growth for a number of yeast species^45^.

### Generating sequence representations and downstream predictions using linear models

To generate LM sequence representations, we pass each sequence to the language models and extract the sequence representation of the last 4 layers for all tokens, which are then mean pooled to obtain a single embedding per sequence as in Devlin et al.^13^ As our models expect fixed length sequences, we truncated longer input sequences. We did this in the following way: For the 3’ sequences, we truncate from the end, for the 5’ sequences, we truncate from the start. If sequences were shorter than the input, for the 3’ model, we feed them as is. For the 5’ model, we pad them from the left using a fixed sequence. We do this because the 5’ model expects the 1000th sequence position to be immediately 5’ of the start codon.

For predicting half-life, we provided 5’ UTR and coding sequence (CDS) features, as described in Cheng et al.^41^ for *S. cerevisiae* and Eser et al.^43^ for *S. pombe*, in addition to our (or competing methods’) sequence representations of the 3’ sequence to a ridge regression with the regularization parameter set by nested cross-validation. We performed 10-fold cross validation to estimate generalization performance.

We proceeded similarly for predicting reporter expression in the experiment of Shalem et al.^40^ but did not include any features besides the 3’ sequence representation. However, as the reporter contains many almost-duplicate sequences which represent mutations of the same endogenous sequence, we performed grouped 10-fold cross-validation, where the groups consisted of said endogenous sequences. This prevented trivial overfitting.

Zrimec et al.^44^ trained different convolutional neural networks using different parts of the regulatory sequence as input. To mimic this, we trained separate linear models for 3’ and 5’ as well as the combination thereof to predict gene expression. Here, we do not perform cross validation but use the same train test split as in Zrimec et al. To predict expression driven by terminator and promoter sequences, Zrimec et al. do not train new models but instead directly apply their convolutional neural network trained on endogenous gene expression. To be comparable, we equally transfer linear model weights to these tasks.

For the microarray, we embed 3’ and 5’ sequences for each species and train separate ridge regressions for each species and condition combination. We again use 10-fold cross validation to evaluate performance.

To assess statistical significance, we computed the residuals and for each task performed a paired Wilcoxon test to determine if the residuals of the Species LM were significantly smaller than those of the next best performing method. The only exception to this is the Zrimec et al. endogenous gene expression task. Here, we only had access to the aggregate performance measures (R^2^) of their models. Thus, we used bootstrapping to compute 95% confidence intervals for the performance of the Species LM and determined its performance to be significantly better than Zrimec et al. if the confidence interval did not overlap their result.

### Modisco Clustering

For de novo motif discovery based on reconstruction probability only, we normalized the reconstruction probability *p* at each position: *p*_*normalized*_ = *p* * *log*(*p*/*p*_*mean*_). We then passed this to tfmodisco-lite (https://github.com/jmschrei/tfmodisco-lite) to obtain motifs.

To perform motif discovery on downstream tasks, we calculated embeddings and predictions for all possible single nucleotide polymorphisms of the original input sequences (in silico mutagenesis). Since ridge models were trained using cross validation, we always select the model which had the non-mutagenized sequence in its test fold to predict mutation effects for this sequence. Having collected all variant effects, we removed their mean at each position (to also associate an attribution score to the reference nucleotide) and passed this to Modisco.

For every Modisco analysis, we set the sliding window size to 8, the flank size to 3, the target seqlet FDR to 0.05 and the number of Leiden runs to 3.

## Acknowledgements

This study was funded by the German Bundesministerium für Bildung und Forschung (BMBF) through the Model Exchange for Regulatory Genomics project MERGE (031L0174A to JG). The funders had no role in the study design, data collection, analysis, decision to publish, or preparation of the manuscript.

We thank all the scientists who isolated, sequenced and assembled the genome data which builds the basis for our research. Figure 1D was created with BioRender.com.

## Model availability

The species and agnostic LMs are available on figshare: https://doi.org/10.6084/m9.figshare.23732655

## Notes

https://doi.org/10.6084/m9.figshare.23732655

### Competing Interest Statement

The authors have declared no competing interest.

### Summary of Updates

The manuscript underwent a major overhaul, providing significantly deeper analyses enabled by more powerful models.

