## Supplementary Figures for "Species-aware DNA language models capture regulatory elements and their evolution"

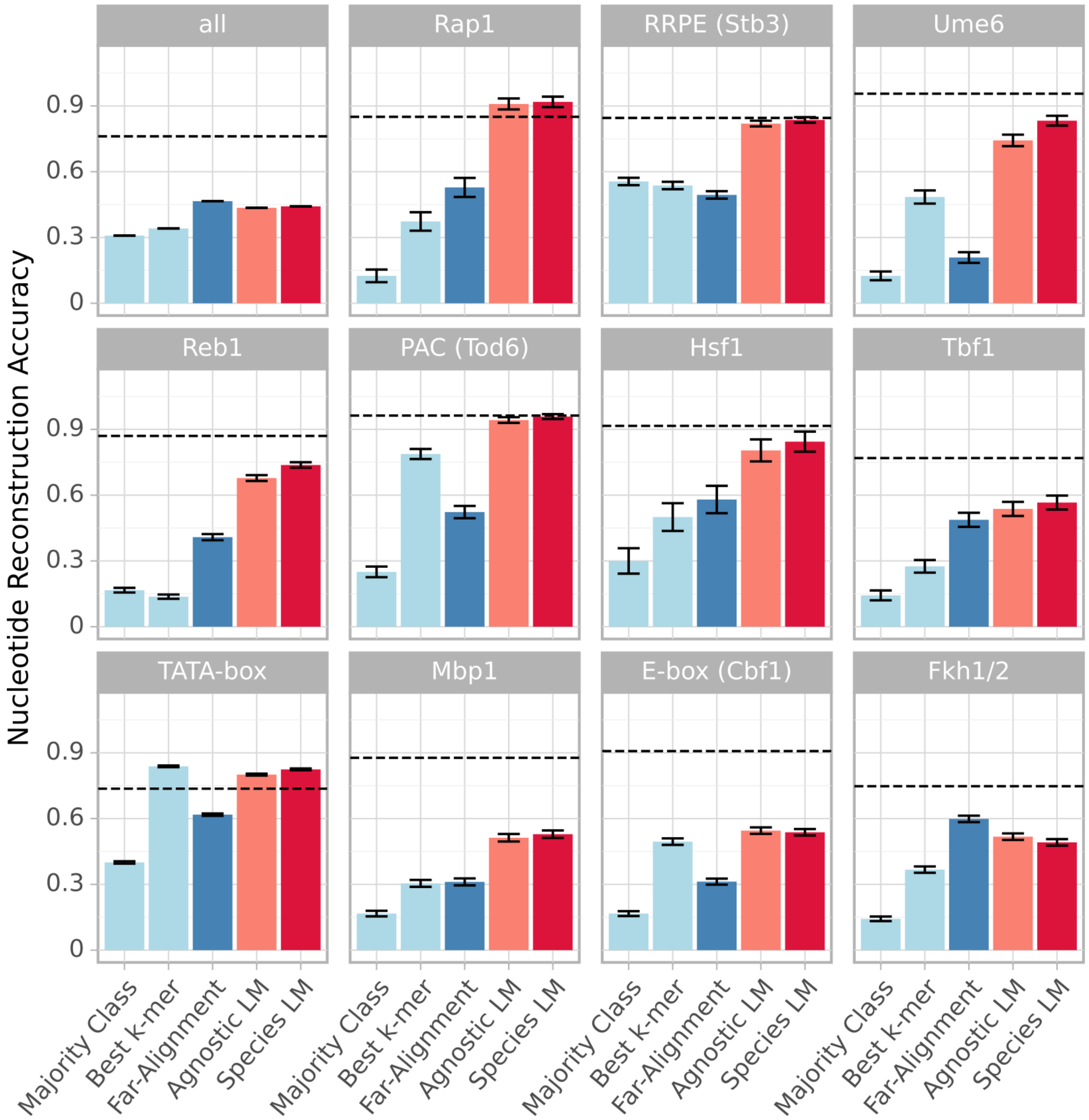


**SFig. 1: Language models reconstruct likely regulatory sequences in the 5' regions of a held-out species.** Reconstruction accuracy for nucleotides within instances of Transcription Factor consensus motifs and across all nucleotides in *S. cerevisiae* 5' regions. We compare the agnostic and species 5' LM to a variety of baselines. Dashed line represents the accuracy achieved by the intra-genus alignment. In many motifs the LMs clearly outperform the baselines. Note that particularly in shorter motifs the reconstruction given here is likely an underestimate, since we consider all matches to the consensus even if many are likely non-functional in-vivo due to their context.

The used consensus motifs are:

- **Rap1**: CAYCCRTACAY
- **RRPE**: AATTTTTCA
- **Ume6**: TAGCCGCC
- **Reb1**: MGGGTAA
- **PAC**: GMGATGAGMT
- **Hsf1**: GAANNTTCTRGAA
- **Tbf1**: ARCCCTAA
- **TATA-box**: TATAWAWR
- **Mbp1**: ACGCGT
- **E-box**: CACGTG
- **Fkh1/2**: GTAAACA

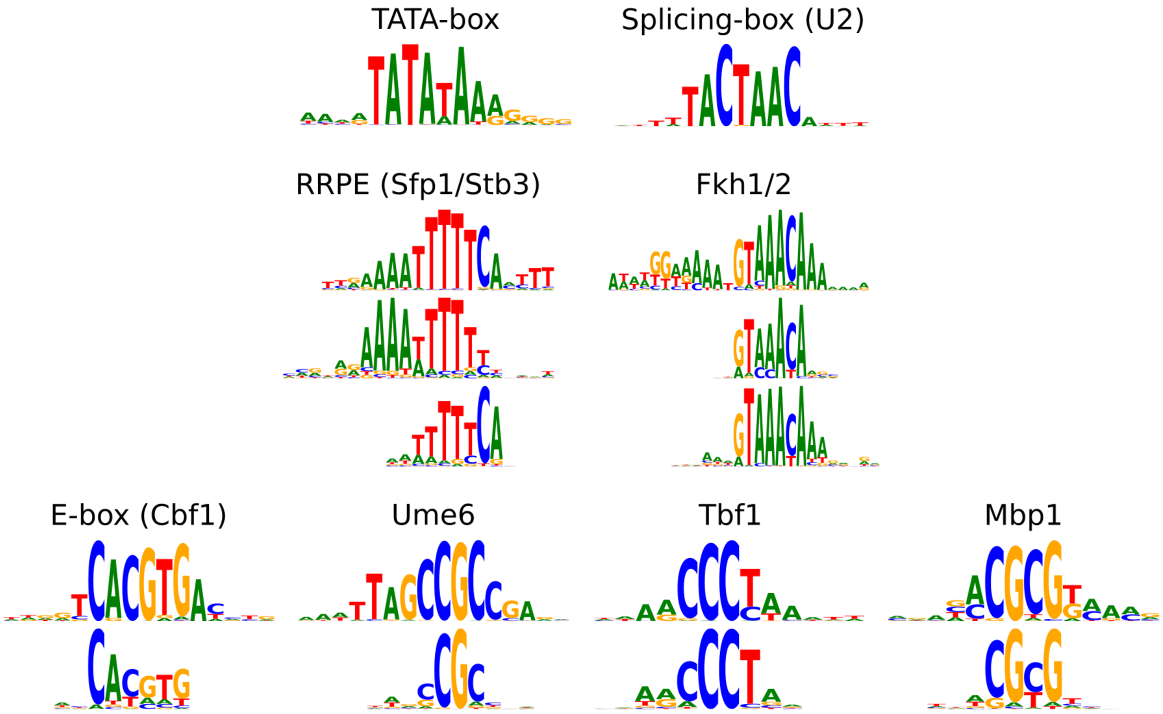


**SFig. 2: Language models recover known binding motifs.** Known transcription factor binding motifs (as well as the splicing motif recognized by the U2-like snRNA) recovered by applying Modisco clustering to the 5' species LM reconstructions. Where possible, the Modisco motif (above) is manually matched to the respective high-confidence PWM from the Yetfasco database (below). We note that the Yetfasco Ume6 motif is derived from in-vitro data, in Chip-exo data the binding preference of Ume6 is much more resemblant of the motif found by Modisco.

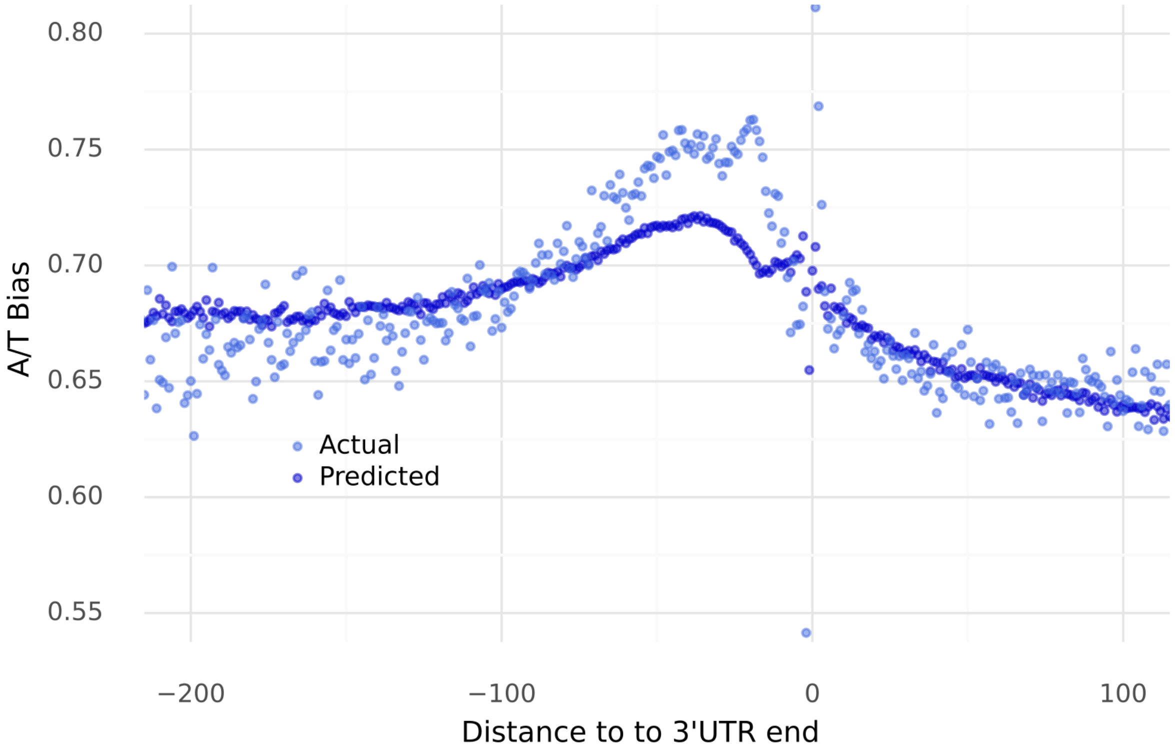


**SFig. 3: The model mimics the elevated AT-bias 5’ of the polyadenylation site.** Actual and predicted (by the 3’ species LM) nucleotide biases as function of the distance to the end of the 3’ UTR. The model keeps track of local variations in AT bias.

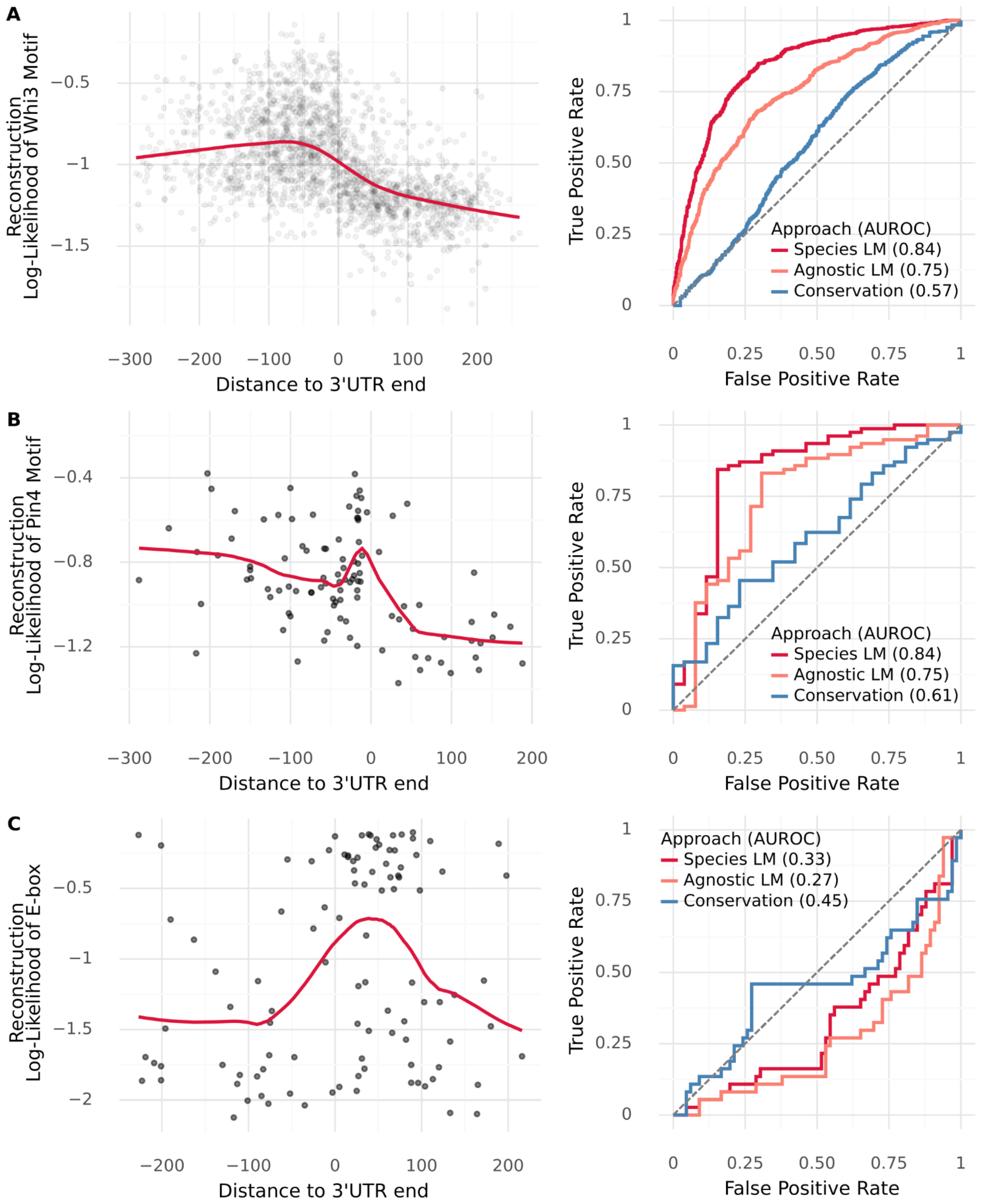


**SFig. 4: Reconstruction of motifs depends on context. A)** Left: Reconstruction fidelity (log-likelihood of the observed nucleotides according to the 3' Species LM) of instances of the Whi3 motif (TGCAT), as function of the distance to the end of the annotated 3'UTR. Right: ROC curve evaluating to what extent the reconstruction fidelity of our 3' LMs, as well as the phastCons conservation score, can serve as a predictor of whether a Puf3 motif instance is within or beyond the 3'UTR boundary. **B)** Same for Pin4 (TTTAATGA). **C)** Same for the E-box (CACGTG), a transcription factor binding motif. We expect TF motifs to be depleted in transcribed regions and indeed the reconstruction fidelity peaks after the end of the 3'UTR.

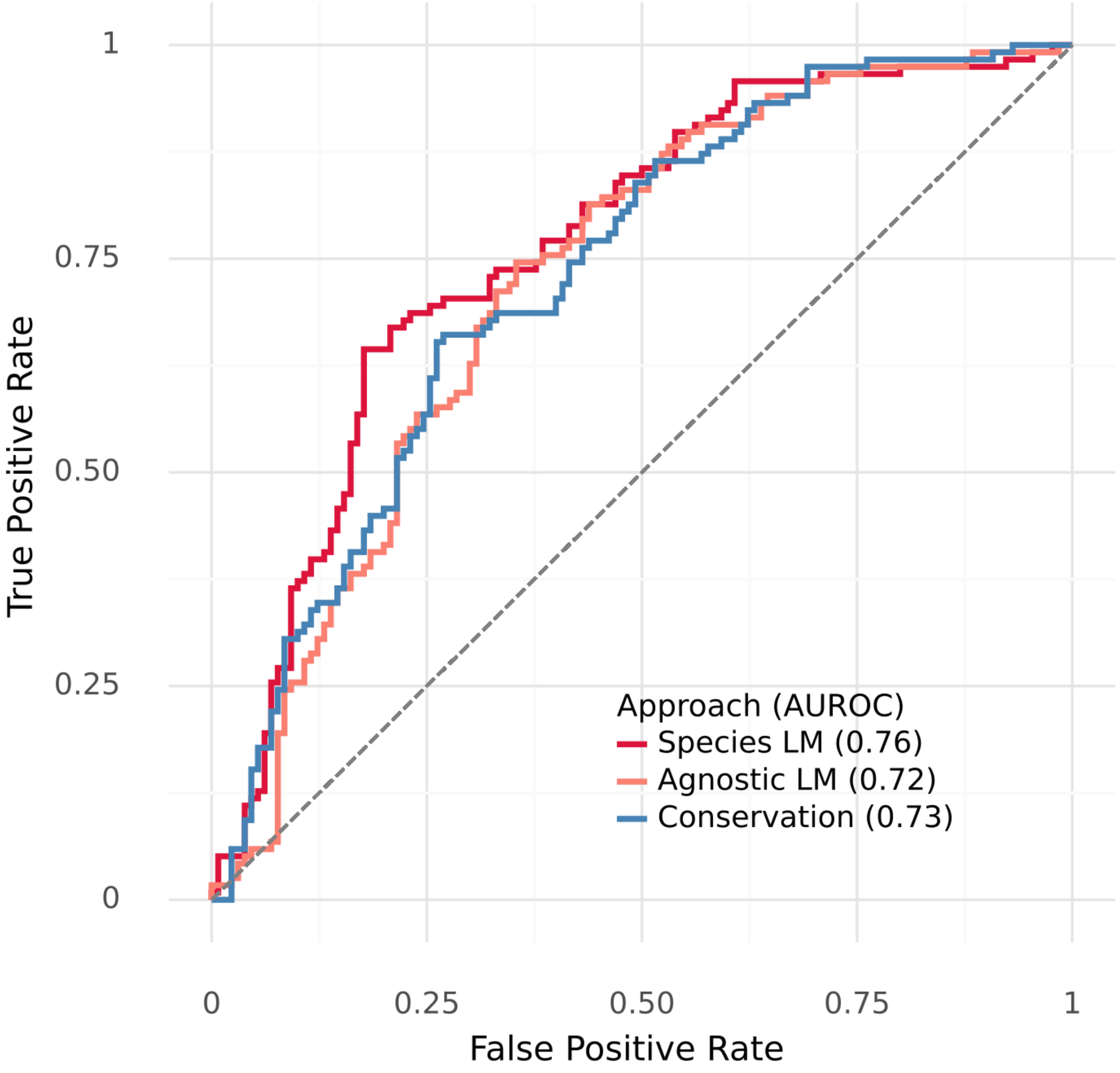


**SFig. 5: Reconstruction fidelity predicts whether a Puf3 motif instance is located 3' of a gene bound by Puf3p in-vivo.** We use the reconstruction fidelity of a Puf3 motif (TGTAAATA) instance achieved by the 3' LMs as a classifier of whether there is experimental evidence that the upstream gene is bound by Puf3p. We also compare against using the phastCons conservation score, which in this case performs on par with the LMs.

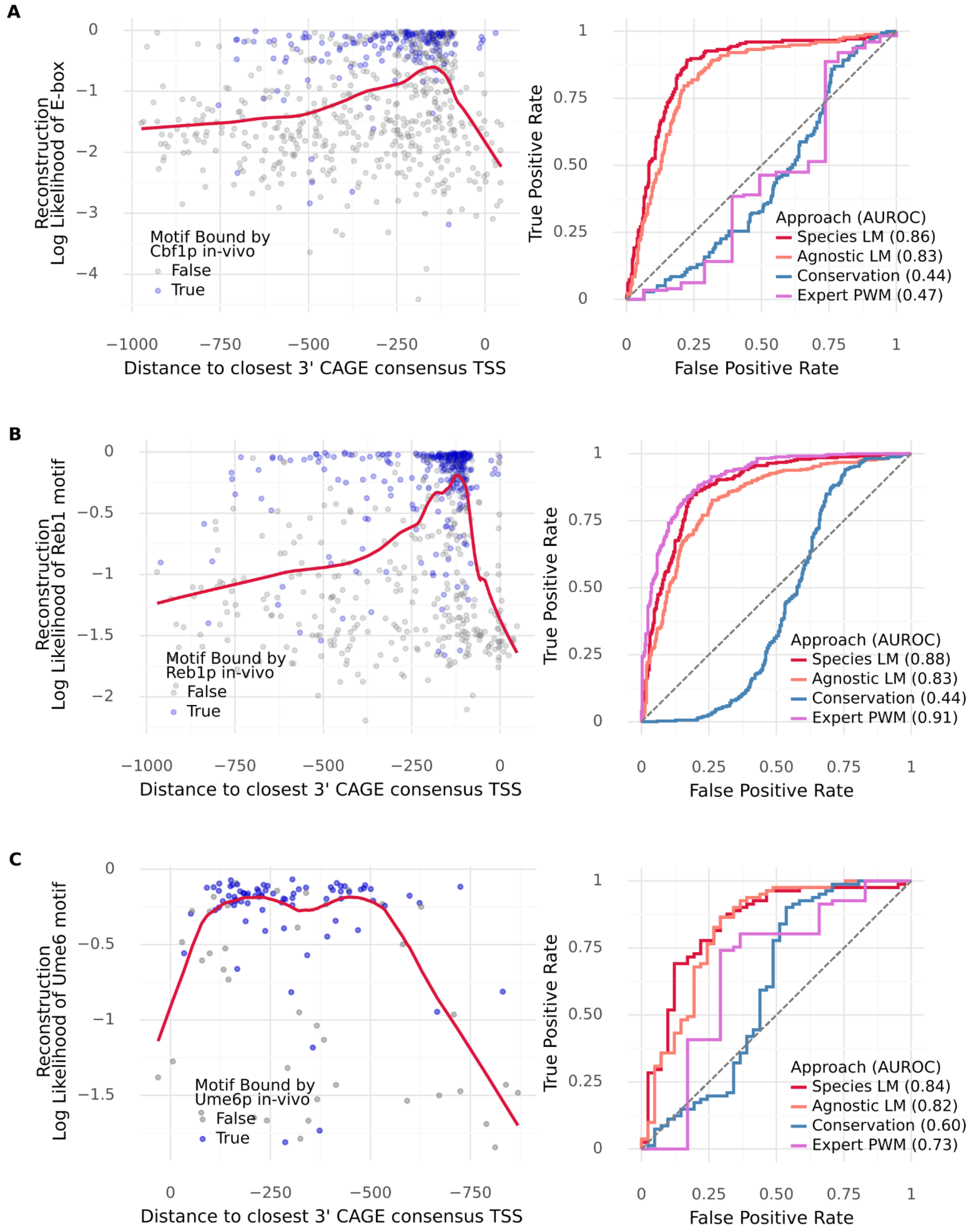


**SFig. 6: Reconstruction of TF motifs depends on context and predicts whether a motif instance will be bound in-vivo. A)** Left: Reconstruction fidelity (log-likelihood of the observed nucleotides according to the 5' Species LM) of instances of the E-box (CACGTG), as function of the distance to the closest 3' TSS (imputed using CAGE data). Blue indicates that the motif instance was bound in-vivo according to Chip-exo data. Right: ROC curve evaluating to what extent the reconstruction fidelity of our 5' LMs, as well as the phastCons conservation score and an expert curated PWM, can serve as a predictor of whether a E-box motif instance is bound in-vivo by Cbf1p. **B)** Same for the Reb1 motif (MGGGTAA). **C)** Same for the Ume6 motif (TAGCCGCC).

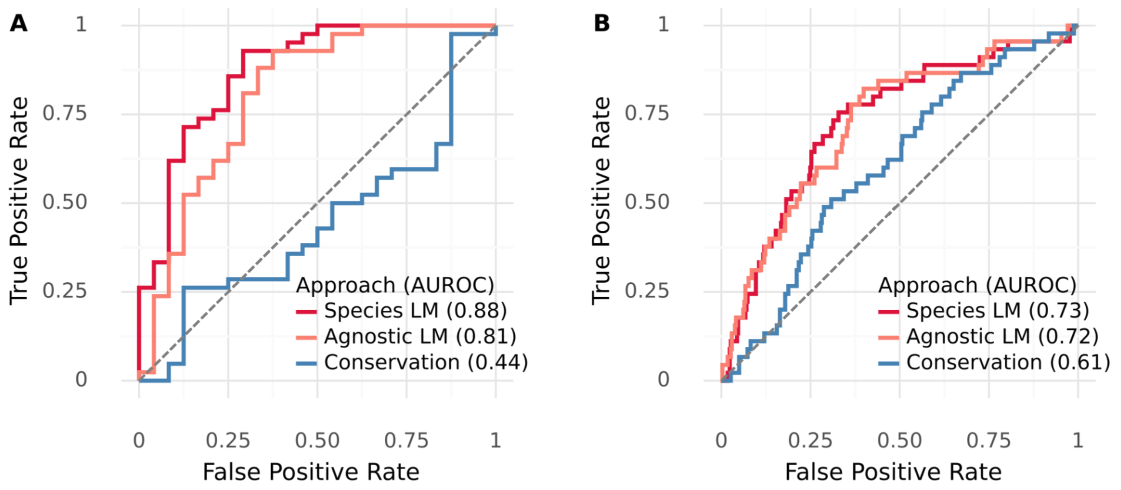


**SFig. 7: Reconstruction fidelity is predictive of the gene module. A)** ROC curve evaluating to what extent the reconstruction fidelity of our 3’ LMs, as well as the phastCons conservation score, can serve as a predictor of whether a Rap1 motif instance (CAYCCRTACAY) is located within 1kb 5’ of a gene which forms part of the ribosomal protein (RP) module. **B)** ROC curve evaluating to what extent the reconstruction fidelity of our 3’ LMs, as well as the phastCons conservation score, can serve as a predictor of whether a RRPE (Stb3) motif instance (AATTTTTCA) is located within 1kb 5’ of a gene which forms part of the ribosome biogenesis module.

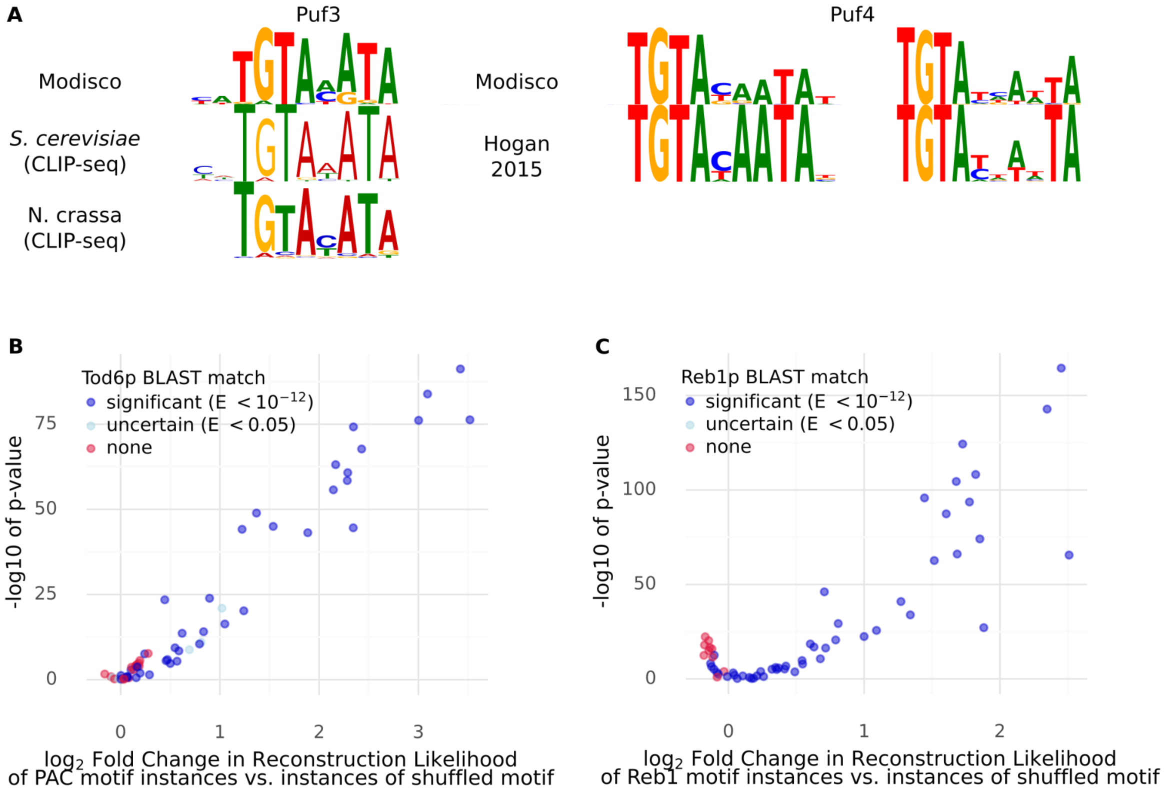


**SFig. 8: LMs can trace the presence and disappearance of motifs across species. A)** Motifs recovered by Modisco clustering to the predictions of the 3' species LM on the 3' regions of all CBP3 homologues in our dataset. We recovered a Puf3 motif and two Puf4 motifs. As we are clustering across species, the recovered Puf3 motif appears to be a hybrid between the *S. cerevisiae* Puf3 motif (which has a preference for a C at -2 and an A in the middle) and the *N. crassa* Puf3 motif (which seemingly has a slight preference for C in the middle). **B)** We computed the reconstruction fidelity (log-likelihood) achieved by the species 5' LM for the *S. cerevisiae* consensus PAC motif instances and for instances matching shuffled versions of this motif in 60 fungal species. The difference in reconstruction between the true and shuffled motif instances, expressed as log2 fold change, is plotted against the -log10 p-value of this difference, computed using a Mann-Whitney-U test. We observe that in species which have no BLAST match to *S. cerevisiae* Tod6p, the reconstruction fidelity of the *S. cerevisiae* Tod6 motif is generally not much better than that of shuffled versions thereof. **C)** Same as B), but for Reb1.

| | *S. cerevisiae* | *S. pombe* | *S. japonicus* | *V. polyspora* | *D. hansenii* | *C. albicans* | *C. glabrata* | *K. lactis* |
|---|---|---|---|---|---|---|---|---|
| Best k-mer | 0.081 | 0.146 | 0.129 | 0.218 | 0.078 | 0.125 | 0.181 | 0.115 |
| Agnostic LM | 0.317 | 0.242 | 0.219 | 0.410 | 0.269 | 0.301 | 0.425 | 0.310 |
| Species LM | **0.366** | **0.341** | **0.286** | **0.437** | **0.321** | **0.327** | **0.456** | **0.342** |

**STable 1: Sequence representations of the species LM outperform other methods on microarray data of Thompson et al.**

Performance ($R^2$) of linear models trained on embeddings from language models compared to best k-mer count regressions, where the best k from {3, 4, 5} is shown.

| Model | *S. cerevisiae* RNA half-life, Cheng et al. | *S. pombe* RNA half-life, Eser et al. | *S. cerevisiae* MPRA, Shalem et al. | *S. cerevisiae* gene expression, Zrimec et al. 3' | *S. cerevisiae* gene expression, Zrimec et al. 5' | *S. cerevisiae* gene expression, Zrimec et al. 3' + 5' | Terminator MPRA, Yamanishi et al. | Promoter MPRA, Keren et al. |
|---|---|---|---|---|---|---|---|---|
| 3-mer | 0.580 | 0.433 | 0.276 | 0.046 | 0.055 | 0.079 | 0.020 | 0.007 |
| 4-mer | 0.569 | 0.450 | 0.278 | 0.040 | 0.076 | 0.094 | 0.029 | 0.062 |
| 5-mer | 0.532 | 0.400 | 0.337 | 0.017 | 0.080 | 0.073 | 0.024 | 0.230 |
| SOTA | 0.586 | 0.422 | | 0.230 | 0.414 | 0.492 | 0.099 | 0.440 |
| Nucleotide Transformer | 0.530 | 0.404 | 0.413 | 0.104 | 0.135 | 0.208 | 0.045 | 0.272 |
| Agnostic LM, yeast only, including sacc. genus | 0.578 | 0.470 | 0.609 | 0.243 | | | | |
| Agnostic LM | 0.578 | 0.479 | 0.532 | 0.240 | 0.365 | 0.438 | 0.071 | 0.352 |
| Agnostic LM, including sacc. genus | 0.580 | 0.476 | 0.530 | 0.276 | | | | |
| Species LM, yeast only, including sacc. genus | 0.583 | 0.471 | 0.577 | 0.206 | | | | |
| Species LM | 0.592 | 0.494 | 0.704 | 0.297 | **0.492** | **0.550** | **0.108** | **0.454** |
| Species LM, including sacc. genus | **0.599** | **0.502** | **0.711** | **0.354** | | | | |

**STable 2: Sequence representations of the species LM outperform other methods on a variety of downstream tasks.**

Performance ($R^2$) of linear models trained on embeddings from language models compared to state-of-the-art models and 3-mer, 4-mer and 5-mer count regressions.
